# Nanotrap Particles Improve Nanopore Sequencing of SARS-CoV-2 and Other Respiratory Viruses

**DOI:** 10.1101/2021.12.08.471814

**Authors:** P Andersen, S Barksdale, RA Barclay, N Smith, J Fernandes, K Besse, D Goldfarb, R Barbero, R Dunlap, T Jones-Roe, R Kelly, S Miao, C Ruhunusiri, A Munns, S Mosavi, L Sanson, D Munns, S Sahoo, O Swahn, K Hull, D White, K Kolb, F Noroozi, J Seelam, A Patnaik, B Lepene

## Abstract

Presented here is a magnetic hydrogel particle enabled workflow for capturing and concentrating SARS-CoV-2 from diagnostic remnant swab samples that significantly improves sequencing results using the Oxford Nanopore Technologies MinION sequencing platform. Our approach utilizes a novel affinity-based magnetic hydrogel particle, circumventing low input sample volumes and allowing for both rapid manual and automated high throughput workflows that are compatible with nanopore sequencing. This approach enhances standard RNA extraction protocols, providing up to 40x improvements in viral mapped reads, and improves sequencing coverage by 20-80% from lower titer diagnostic remnant samples. Furthermore, we demonstrate that this approach works for contrived influenza virus and respiratory syncytial virus samples, suggesting that it can be used to identify and improve sequencing results of multiple viruses in VTM samples. These methods can be performed manually or on a KingFisher Apex system.

## Introduction

As of Oct 28, 2021, there have been more than 246 million COVID-19 cases and nearly 5 million COVID-19 related deaths worldwide ^1^. Viral mutations have enabled the pandemic to continue pervading everyday life despite the use of widespread global health measures to prevent the spread of SARS-CoV-2. Detection and monitoring of emerging viral variants have become a critical tool in the global health response, highlighting the need for rapidly deployable and accurate sequencing methods ^2–5^. Recent advances in next-generation sequencing (NGS) technologies have made the routine use of sequencing for monitoring and identifying viral outbreaks more possible, but many NGS instruments are not portable and still cost prohibitive, thus limiting their overall adoption ^6,7^. The Oxford Nanopore Technologies (ONT) MinION platform offers a relatively inexpensive and portable detection strategy, one that is capable of identifying and sequencing various respiratory viruses in the field ^8–10^.

While nanopore sequencer advances make rapid on-site detection and characterization of SARS-CoV-2 and other viral genomes a possibility, these portable sequencers are still limited by certain disadvantages, namely that unless large amounts of viral RNA are used for the sequencing reactions, there can be accuracy issues during basecalling ^11–16^. These technical limitations reduce the usefulness of a tool that could improve the ability to rapidly and accurately respond to viral outbreaks and track transmission in real time.

Increasing the total amount of RNA material for analysis through sample enrichment is one strategy available for improving the performance of sequencing platforms. To this end, we sought to address the viral sequencing limitations of a nanopore sequencer by applying the affinity-based magnetic hydrogel particle (Nanotrap particle) enrichment technology to SARS-CoV-2 viral transport medium (VTM) samples. The Nanotrap particle technology has shown broad application in clinical diagnostics by enriching and stabilizing biomarkers and analytes in complex clinical samples. Recent studies demonstrated that Nanotrap particles are able to concentrate and improve detection of many viral types, including SARS-CoV-2, on multiple molecular assays ^17–21^.

When utilizing a portable sequencing platform such as the ONT MinION sequencer, increasing the amount of input RNA should enable successful sequencing of lower titer viral samples, which, in turn potentially increases the fraction of patient samples that are viable for sequencing. Ideally this should improve identification of specific viral mutations, hastening the response to emerging problematic variants ^22–24^.

Here, we show Nanotrap particles can improve current sequencing workflows and enable new ones by enhancing current standard RNA extraction methods. We demonstrate that these Nanotrap particle workflows improve sequencing results by increasing total viral mapped reads, resulting in greater sequencing depth and coverage. Nanotrap particle workflows were developed for multiple RNA extraction kits, and their utility is demonstrated in both contrived and diagnostic remnant samples.

## Methods and Materials

### Magnetic Hydrogel Particles

Nanotrap® Magnetic Virus Particles (SKU: 44202) were provided by Ceres Nanosciences Inc. Manassas, VA.

### Biological Materials

Contrived samples were comprised of heat inactivated virus spiked into VTM (Puritan UniTranz-RT Transport Systems, cat# 89233-458). Heat-inactivated viruses were purchased from Zeptometrix: SARS-CoV-2 (cat# 0810587CFHI), Influenza A-H1N1 (cat# 0810109CFHI), and Respiratory Syncytial Virus Type A(RSV) (cat# 0810040ACFHI)

### Clinical Samples

SARS-CoV-2 positive diagnostic remnant samples in viral transport medium were purchased from Discovery Life Sciences, Huntsville, AL. Discovery Life Sciences previously tested these samples by RT-PCR, obtaining cycle thresholds ranging from 24 to 35.

### Nanotrap Virus Capture Workflows

#### Nanotrap Particle Workflow 1

(Manual Method with Column-Based RNA Extraction Kit): Two-hundred microliters of Nanotrap particles were added to 1,000 microliters VTM samples. Samples were incubated at room temperature for 10 minutes, and then placed on a magnetic separator for 1-2 min to allow the Nanotrap particles to pellet. Supernatants were removed and discarded. One hundred microliters of RNAse/DNase-free water with 350 microliters of QIAGEN Buffer RLT were added to the Nanotrap particle pellet. Samples were incubated for 10 min on a shaker at room temperature before being placed on a magnetic separator for 1-2 min to allow the Nanotrap particles to pellet. Supernatants containing the viral nucleic acid material were processed for RNA extraction using the QIAGEN RNeasy MinElute Cleanup Kit (cat #74204) following the manufacturer’s 100 μL workflow instructions. Following RNA extraction, RNA samples were ready for sequencing library preparation.

#### Nanotrap Particle Workflow 2

(Manual Method with Magnetic Bead-Based RNA Extraction Kit): Three hundred microliters of PBS with 0.05% Tween-20 (v/v) was added directly to 500 microliters VTM samples. Two hundred microliters of Nanotrap particles were added to each VTM sample, which were then incubated at room temperature for 10 minutes. Samples were placed on a magnetic separator for 1-2 min to allow the Nanotrap particles to pellet. Supernatants were removed and discarded. Nanotrap particle pellets were resuspended in 1000 microliters of molecular grade water with 0.05% Tween-20 (v/v). Following a brief resuspension, the samples were again placed on a magnetic separator for 1-2 min to allow the Nanotrap particles to pellet, and the supernatant removed and discarded. Nanotrap particles were resuspended in 200 microliters of MagMAX Microbiome Lysis Solution, and samples were incubated at 65°C on a shaker for 10 minutes. Samples were placed on a magnetic separator to allow the Nanotrap particles to pellet for 1-2 min. Supernatants containing the viral nucleic acid material underwent RNA extraction using the using the MagMAX Microbiome Ultra Nucleic Acid Isolation Kit (cat# A42357) following the manufacturer’s 200 microliters workflow instructions. Following extraction kit processing, RNA samples were ready for sequencing library preparation.

#### Nanotrap Particle Workflow 3

(Automated Method with Magnetic Bead-Based RNA Extraction Kit): The following method used the KingFisher Apex System and associated consumables. Three hundred microliters of PBS with 0.05% Tween-20(v/v) was added directly to 500 microliters VTM samples in a 96 deep well KingFisher plate. Two-hundred microliters of Nanotrap particles were added to each VTM sample. Molecular grade water with 0.05% Tween-20 (v/v) was added to a second 96 DW plate, and 200 microliters of MagMAX Microbiome Lysis Solution was added to a third 96 DW plate. A custom KingFisher program “NT2MM.kfx” was made to process the Nanotrap particles using the three prepared 96 DW plates. The entire process occurred in 30 min, with the final eluate containing extracted viral RNA.

After Nanotrap particle processing, extracted viral RNA samples were processed using the MagMAX Microbiome Ultra Nucleic Acid Isolation Kit following the manufacturer’s 200 microliters workflow instructions for use with KingFisher Apex. Following extraction kit processing, RNA samples were ready for sequencing library preparation.

### RNA Extraction Kits

The Nanotrap particle workflows described above were benchmarked against the QIAGEN and ThermoFisher MagMAX RNA extraction kit workflows without any Nanotrap particles. For Workflow 1 comparison, samples were processed using the QIAGEN RNeasy MinElute Cleanup Kit(cat #74204) following the manufacturer’s 100 microliters workflow instructions. For Workflow 2 and 3 comparisons, samples were processed using the ThermoFisher MagMAX Microbiome Ultra Nucleic Acid Isolation Kit (cat# A42357) following the manufacturer’s 200 microliters workflow instructions.

### RT-PCR

For RT-PCR analysis of SARS-CoV-2 samples, the IDT 2019 nCoV CDC EUA Kit (cat# 1006770), which includes N1 primers/probes, was used for real-time RT-PCR. Following IDT’s recommendation, TaqPath 1-Step RT-qPCR Master Mix from ThermoFisher (cat# A15300) was used in the IDT 2019 nCoV CDC EUA assay. Each PCR reaction used 8.5 microliters of nuclease free water, 5 microliters of the TaqPath solution, 1.5 microliters of the N1 primer/probe, and 5 microliters of RNA template. PCR conditions were performed according to IDT’s instructions on a Roche LightCycler 96. All SARS-CoV-2 (both heat-inactivated and diagnostic remnant) experiments utilized this assay.

For RT-PCR analysis of Influenza A and RSV samples, the Primerdesign Influenza A H1 Kit (Path-H1N1-v2.0-Standard) and the Primerdesign RSV kit (Path-RSV-A-Standard) were used following the manufacturer’s instructions. PCR conditions were performed according to Primerdesign’s instructions on a Roche LightCycler 96.

### Library Preparation and Sequencing Workflow

After viral RNA extraction, samples were prepared for sequencing using the ARTIC Network developed; “nCoV-2019 Sequencing Protocol v3(Lo Cost)” ^25^. Briefly summarized, amplified cDNA was prepared using a targeted amplicon approach. Per the ARTIC nCov-2019 protocol, IDT ARTIC V3 Amplicon Sequencing Panel primers were used. These 218 primers, covering the entire SARS-CoV-2 genome, were used to generate and amplify cDNA from the extracted viral RNA. Once cDNA was prepared, the samples were processed using the Oxford Nanopore Technologies (ONT) Ligation Sequencing Kit, barcoded individually using the ONT Native barcoding expansion kit native barcodes with a modified “One-pot” protocol. These individual samples were pooled together and concentrated using AMPure XP magnetic beads (cat# A63880) following the nCoV-2019 protocol modifications. The pooled library was loaded onto an ONT FLO-MIN106 R.9 flow cell used with the ONT Mk1C Sequencing Platform. Unless otherwise stated, the ONT Mk1C was run for 24 hrs using the LSK109 kit with EXP-ND-196 barcodes selected.

### Bioinformatics and Data Analysis

To analyze and process the sequencing data generated by the ONT Mk1C platform, the following tools were used: live basecalling and demultiplexing was performed using the ONT MinKnow software integrated into the ONT Mk1C MinION device; general classification and viral mapped reads were generated using the 3/9/2020 W.I.M.P protocol through ONT’s Epi2me web tool; further coverage analysis was conducted using Minimap2 and Samtools through the UseGalaxy.org web portal. Statistical analyses were performed and figures were generated using Graphpad Prism 9.

## Results

Prior studies have demonstrated that Nanotrap particles capture and concentrate multiple respiratory viral pathogens, including SARS-CoV-2, Influenza A, Influenza B, and RSV ^17–21^. This enrichment led to improved results by various molecular assays, including real-time PCR. We hypothesized that if the Nanotrap particles could improve those molecular assays, they would also improve sequencing platforms by increasing the amount of viral RNA available for sequencing. To demonstrate the robustness and ease-of-use of the Nanotrap particle technology, three workflows were developed - a manual method with a column-based RNA extraction kit, a manual method with a magnetic-bead-based RNA extraction kit, and an automated method with a magnetic-bead-based RNA extraction kit.

### Nanotrap Particles Improve ONT Sequencing Results for Contrived SARS-CoV-2 VTM Samples using a Column-Based RNA Extraction Method

We developed a method utilizing Nanotrap particles to capture and concentrate virus followed by a column-based RNA extraction, examining the particles ability to improve nanopore sequencing of SARS-CoV-2. To that end, neat VTM samples were spiked with 1:10 serial dilutions of heat-inactivated SARS-CoV-2 starting from 10^6^ TCID_50_/mL to 10^2^ TCID_50_/mL. Samples were processed with Nanotrap Particle Workflow 1 using the QIAGEN RNeasy MinElute Cleanup Kit for the viral RNA extraction ([+NT]) or without Nanotrap particles using the RNeasy kit alone ([−NT]). The extracted RNA samples were prepared for sequencing using the ARTIC nCoV-2019 sequencing protocol and run on the ONT Mk1C sequencer as described in the method section. Extracted viral RNA was also analyzed using the IDT 2019 nCoV CDC EUA RT-PCR Kit to identify a potential correlation between the two assays and confirm the presence of SARS-Cov-2. As shown in **Figure 1A**, Nanotrap particles improved sequencing results at multiple concentrations when compared to the workflow without Nanotrap particles. A 6.0x improvement in SARS-CoV-2 viral mapped reads was observed at 10^6^ TCID_50_/mL and a 2.0x improvement was seen at 10^5^ TCID_50_/mL. Statistical analysis showed improvements were significant with p-values of <0.05 for both concentrations of virus. No significant improvement was seen between the [+NT] and [−NT] samples below 10^5^ TCID_50_/mL. When RT-PCR was performed, Nanotrap particles improved viral recovery by 2 PCR Cycle thresholds (Cts) across the first four serial dilutions (**Figure 1B**). Paired t-tests confirmed significant improvement for the same four concentrations.

**Fig. 1:**
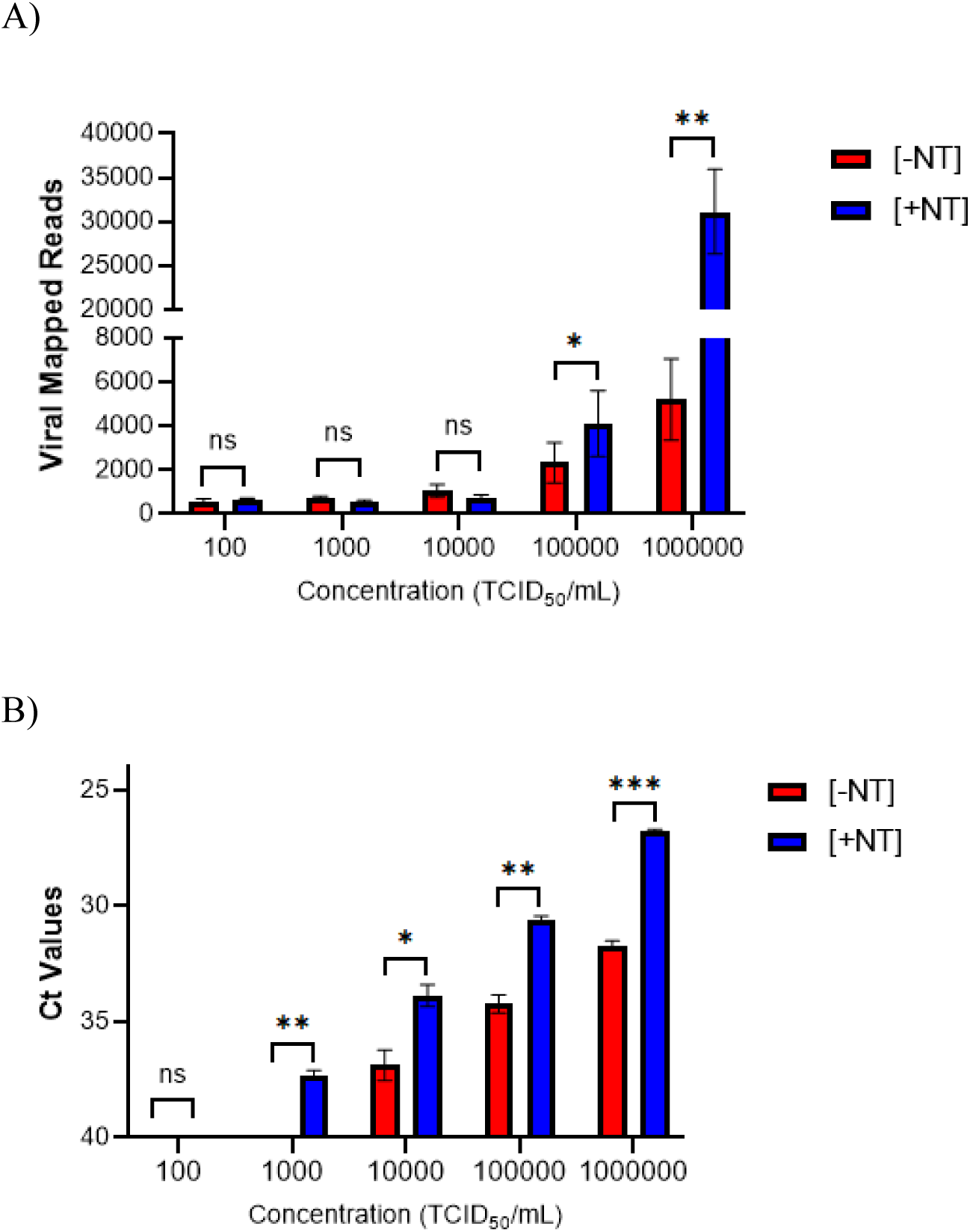
Nanotrap Particle Workflow 1 Improves Sequencing of Contrived SARS-CoV-2 Samples. Heat inactivated SARS-CoV-2 was spiked into VTM at 10^2^, 10^3^, 10^4^, 10^5^, and 10^6^ TCID_50_/mL and samples were processed using Nanotrap Particle Workflow 1 [+NT] or the RNEasy Kit alone [−NT]; n=3 for each process. Samples then underwent sequencing on a ONT MinION R.9 flow cell (**A**) or RT-PCR (**B**). [+NT] were compared to [−NT] by paired *t*-test in order to assess significance of increased viral detection. * p<0.05, ** p<0.01, *** p<0.001.

### Nanotrap Particles Improve ONT Sequencing Results for Contrived SARS-CoV-2 VTM Samples Using a Magnetic-Bead-Based RNA Extraction Kit

Column RNA extractions are typically used in low-sample-throughput, high-complexity laboratory benchtop environments. Given these limitations, we assessed the Nanotrap particles’ ability to improve an RNA extraction method based on magnetic beads. Nanotrap Particle Workflow 2 was tested in a similar manner to Workflow 1: neat VTM samples were spiked with a 1:10 serial dilution of SARS-CoV-2 starting from 10^6^ TCID_50_/mL down to 10^2^ TCID_50_/mL. Samples were processed with ([+NT]) or without ([−NT]) Nanotrap particles. The [−NT] sample was processed using the MagMAX Microbiome Ultra Nucleic Acid Isolation Kit alone.

Following RNA extraction, samples were then prepared for sequencing using the ARTIC nCoV-2019 Sequencing protocol, run on the ONT Mk1C Sequencing Platform. Processed RNA samples were also analyzed by RT-PCR using the IDT 2019 nCoV CDC EUA RT-PCR Kit to confirm the presence of virus.

Nanotrap particles improved sequencing results at multiple concentrations when compared to the [−NT] workflow (**Figure 2A**). A 1.9x improvement in SARS-CoV-2 viral mapped reads was observed at 10^6^ TCID_50_/mL and a 1.4x improvement was seen at 10^5^ TCID_50_/mL. Statistical analysis confirmed significance, p values of <0.05 were calculated for both results. Additionally, Nanotrap particles improved SARS-CoV-2 detection in RT-PCR, providing an average 1.5 Ct improvement at 10^6^-10^2^ TCID_50_/mL (**Figure 2B**).

**Fig. 2:**
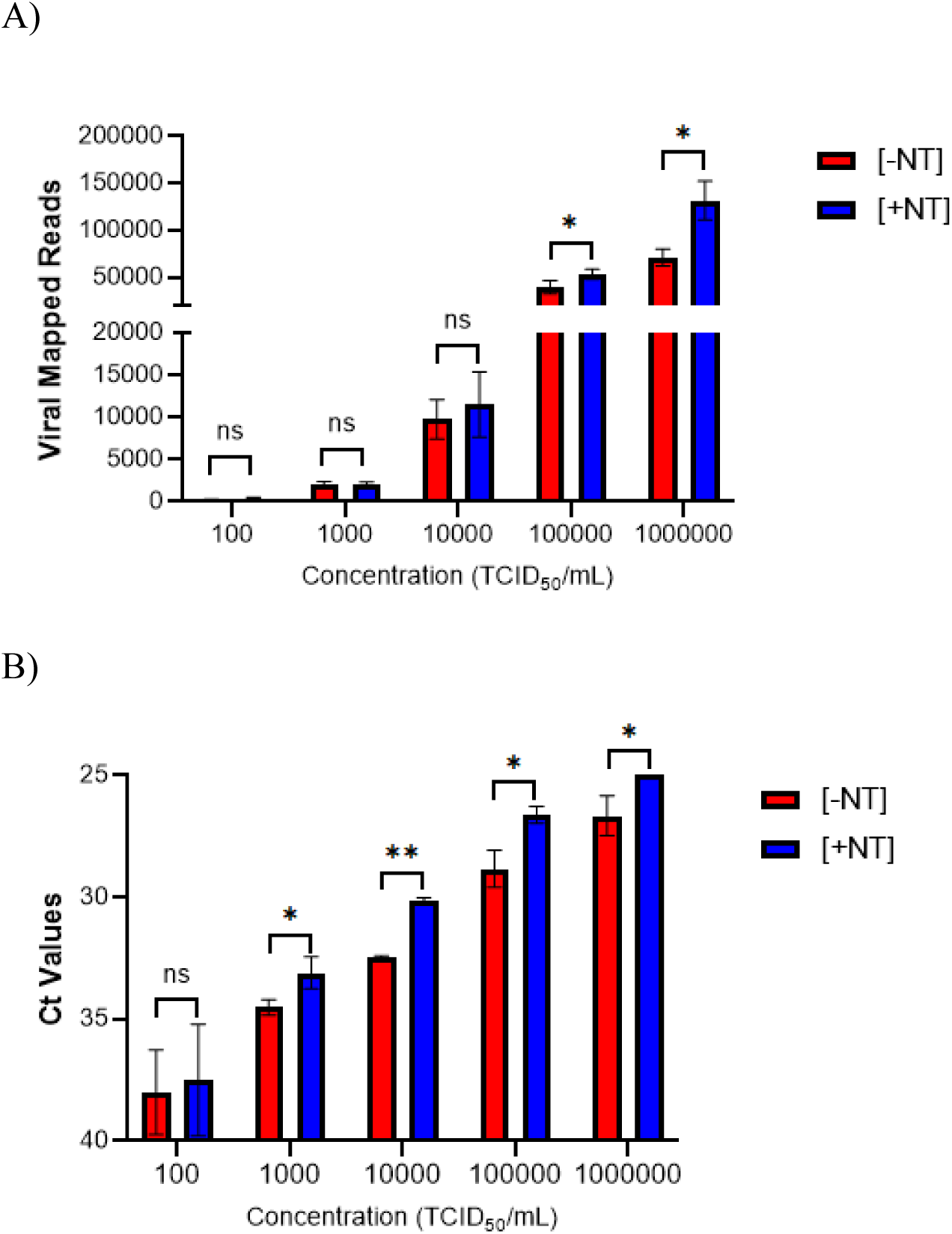
Nanotrap Particle Workflow 2 Improves Sequencing of Contrived SARS-CoV-2 Samples. Heat inactivated SARS-CoV-2 was spiked into VTM at 10^2^, 10^3^, 10^4^, 10^5^, and 10^6^ TCID_50_/mL and samples were processed using Nanotrap Particle Workflow 2 [+NT] or the MagMAX Kit alone [−NT]; n=3 for each process. Samples then underwent sequencing on a ONT MinION R.9 flow cell (**A**) or RT-PCR (**B**). [+NT] were compared to [−NT] by paired *t*-test in order to assess significance of increased viral detection. * p<0.05, ** p<0.01.

### Nanotrap Particles Improve ONT Sequencing Results for Diagnostic Remnant SARS-CoV-2 VTM Samples Using a Column-Based RNA Extraction Method

Contrived VTM samples are useful for evaluating methods in a pristine environment, but they are not necessarily indicative of how a method will work with clinical samples. Thus, we evaluated the Nanotrap particle workflows using diagnostic remnant clinical swab VTM samples. We utilized ten diagnostic remnant VTM specimens with reported RT-PCR test cycle thresholds ranging from 24 to 35 (as reported by the specimen supplier). The same processing and sequencing workflow established with contrived samples was used for these diagnostic remnant samples. To better assess the impact of the Nanotrap particle workflow on sequencing results, we quantified both total viral mapped reads and the associated percent genome coverage at 30x depth of the processed samples. Results in **Figure 3A**, which were generated using Nanotrap Particle Workflow 1[+NT], show that Nanotrap particles improved the sequencing results of 100 % of the diagnostic remnant samples (n = 10). Compared to the workflow without Nanotrap particles[−NT], the use of Nanotrap Particle Workflow 1 resulted in an average 7x improvement in total viral mapped reads across all diagnostic remnant samples. These viral mapped read improvements resulted in an average viral genome coverage increase of 52% over samples processed without Nanotrap particles (**Figure 3B**). A paired t-test across all 10 samples shows the increases are statistically significant for both the viral mapped reads and coverage percent (**Figure 3D, Figure 3E**). In Figure 3C, RT-PCR confirmed the presence of SARS-CoV-2 for all 10 samples, also resulting in an average improvement of 4 Ct over [−NT] samples. It is worth noting that 3 samples were below the detection limit of the RT-PCR assay when processed without Nanotrap particles but that all 10 samples had detectable RNA when the Nanotrap particles were used in sample processing.

**Fig. 3:**
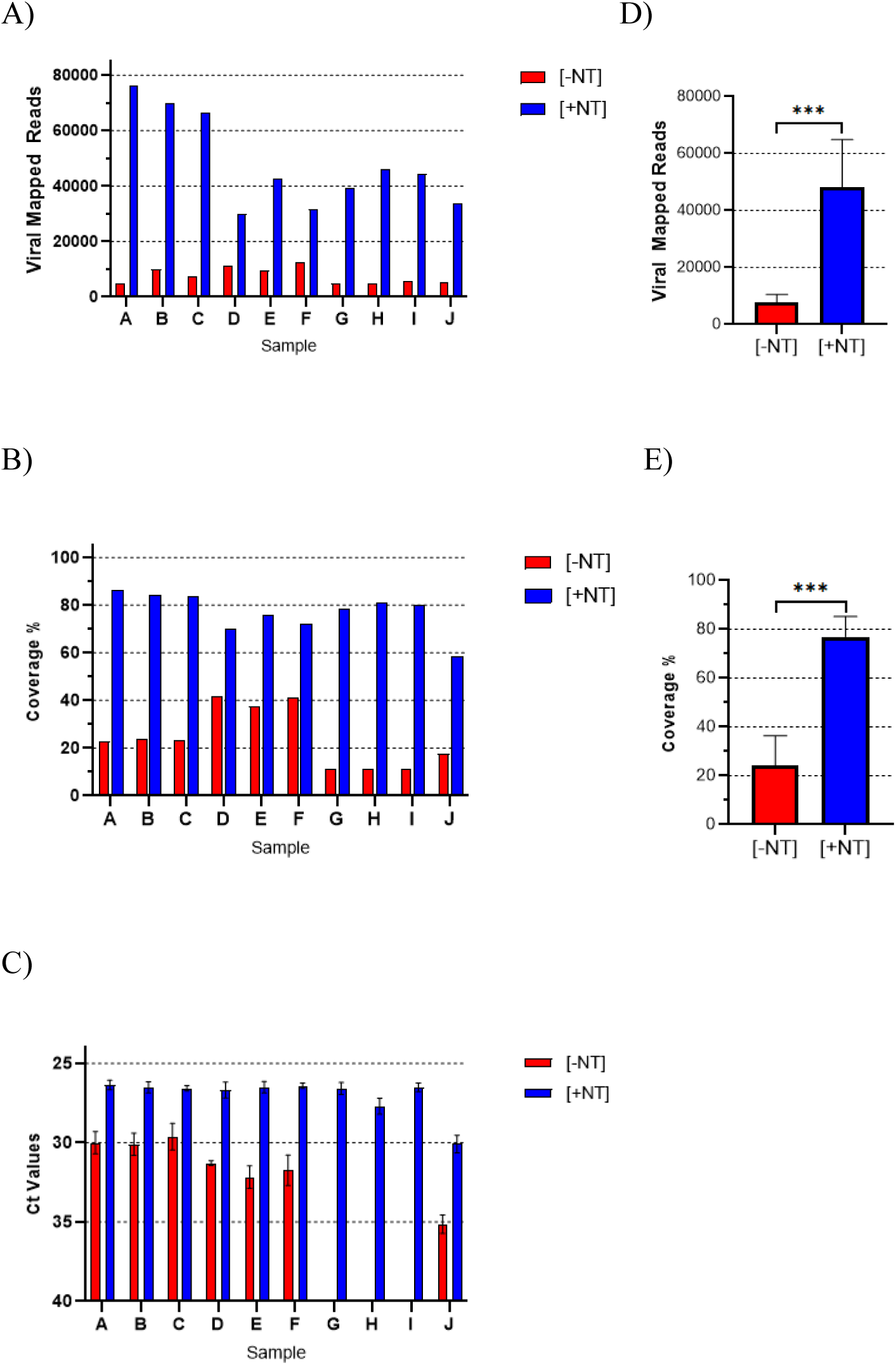
Nanotrap Particle Workflow 1 Improves Sequencing of Diagnostic Remnant SARS-CoV-2 Samples. 10 SARS-CoV-2 positive diagnostic remnant samples were processed using Nanotrap Particle Workflow 1 [+NT] or the RNEasy Kit alone [−NT]. Samples then underwent sequencing on a ONT MinION R.9 flow cell and were analyzed by Viral Mapped Reads to SARS-CoV-2 (**A**), Viral Genome Coverage at 30x depth (**B**), or RT-PCR (**C**). [+NT] were compared to [−NT] by paired *t*-test in order to assess significance of increased viral detection (**D**),(**E**). *** p<0.001.

#### Nanotrap Particles Improves ONT sequencing results for Diagnostic remnant SARS-CoV-2 VTM samples using Using a Magnetic-Bead-Based RNA Extraction Kit on a KingFisher system

One of the advantages of magnetic particle based sample processing is that the method can be readily automated. To that end, we developed an automated version of Nanotrap Particle Workflow 2, to be used on the Kingfisher Apex platform. We then compared this automated Nanotrap Particle Workflow 3 method ([+NT]) to a method without Nanotrap particles using ten additional SARS-CoV-2 positive diagnostic remnant samples ([−NT]), once again examining the sequencing and RT-PCR output of the two methods. We observed that the Nanotrap particle processing significantly improved sequencing results for 7 of 10 samples, resulting in an average improvement of 42x in total viral mapped reads (**Figure 4A**). This corresponded to an average 51% increase in viral genome coverage relative to [−NT] samples (**Figure 4B**). Paired t-tests confirmed that Nanotrap particles significantly improved both viral mapped reads (Figure 4D) and genome coverage (**Figure 4E**). As with the previous set of diagnostic remnant samples, RT-PCR confirmed the presence of SARS-CoV-2 for all 10 samples. The [+NT] automated process improved RT-PCR results as well, resulting in an average 3.7 Ct improvement shown in **Figure 4C**.

**Fig. 4:**
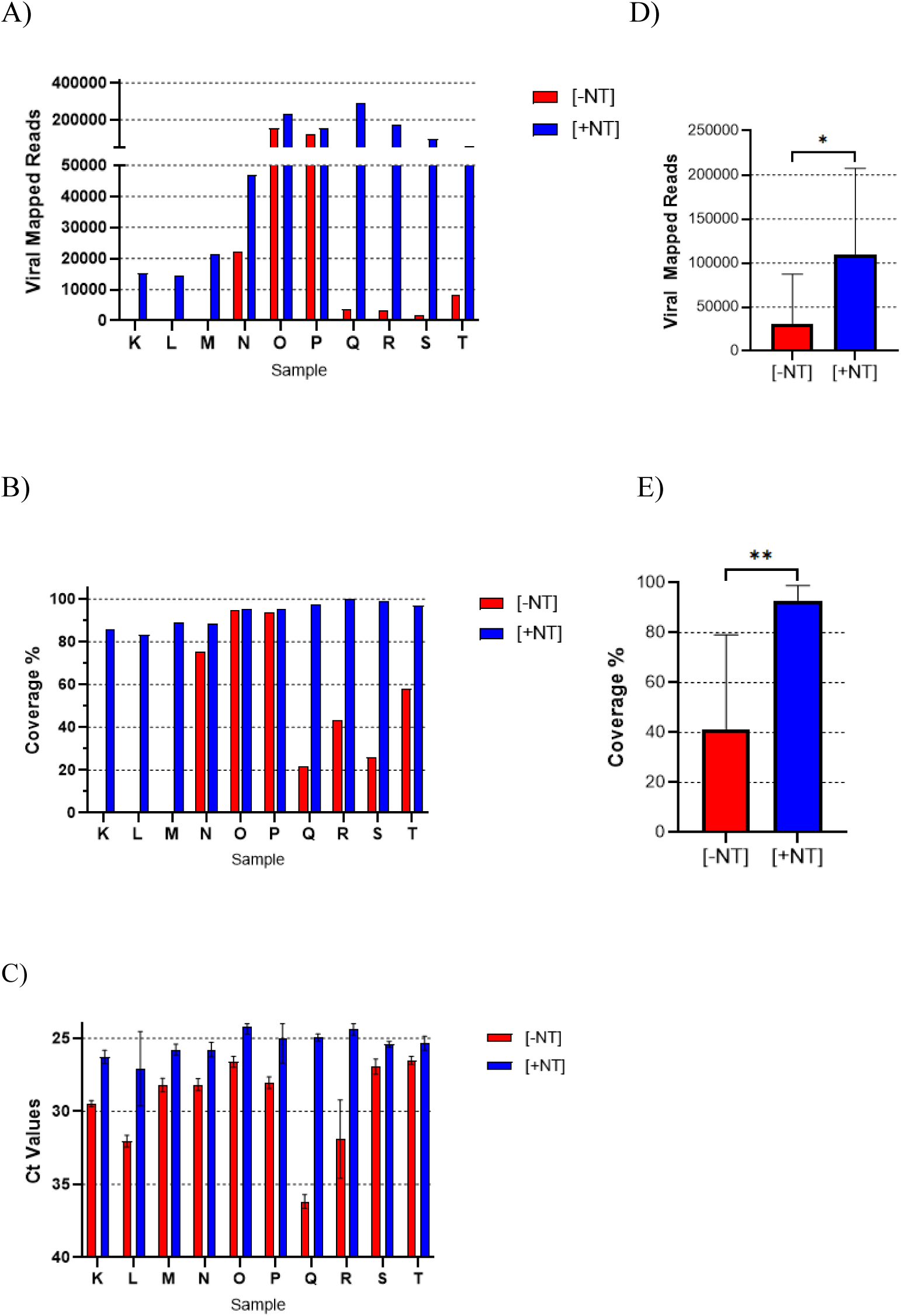
Nanotrap Particle Workflow 3 Improves Sequencing of Diagnostic Remnant SARS-CoV-2 Samples. 10 SARS-CoV-2 positive diagnostic remnant samples were processed using Nanotrap Particle Workflow 3 [+NT] or the MagMAX kit alone [−NT]. Samples then underwent sequencing on a ONT MinION R.9 flow cell and were analyzed by Viral Mapped Reads to SARS-CoV-2 (**A**), Viral Genome Coverage at 30x depth (**B**), or RT-PCR (**C**). [+NT] were compared to [−NT] by paired *t*-test in order to assess significance of increased viral detection (**D**), (**E**). * p<0.05, ** p<0.01.

#### Nanotrap® particles coupled with column-based RNA extraction workflow yields detection of multiple virus types

Ideally, viral concentration technologies should allow for the concentration of multiple viruses, not just SARS-CoV-2. As prior studies have demonstrated Nanotrap particles capture a variety of respiratory viruses, we briefly investigated whether the Nanotrap particles also could be used to improve sequencing of Influenza A (H1N1) and Respiratory Syncytial Virus (RSV). Neat VTM was spiked separately at 10^6^ TCID_50_/mL with inactivated Influenza A and RSV. Using the previously established column based RNA extraction protocol, viral contrived samples were processed with Nanotrap Particle Workflow 1[+NT] comparing the results against samples processed without Nanotraps[−NT]. The resulting RNA eluates were then prepared for sequencing using a modified version of the ARTIC Library Prep protocol using primers specific to Influenza A and RSV. Samples were run on the ONT Mk1C. Results in **Figure 5A** and **Figure 6A** demonstrate that Nanotrap particles improve nanopore sequencing results for contrived Influenza A and RSV samples. Similarly to SARS-CoV-2, Nanotrap particles increased viral mapped reads by 4x for both Influenza A and RSV compared to the samples processed without Nanotrap particles. Additionally, using Nanotrap particles improved the detection of viral RNA as measured by RT-PCR (**Figure 5B, Figure 6B**), paired t-test confirmed the improvement was again statistically significant with calculated p values < 0.05. These results indicate that Nanotrap particles can be used to identify and improve sequencing results of multiple viruses in VTM samples.

**Fig. 5:**
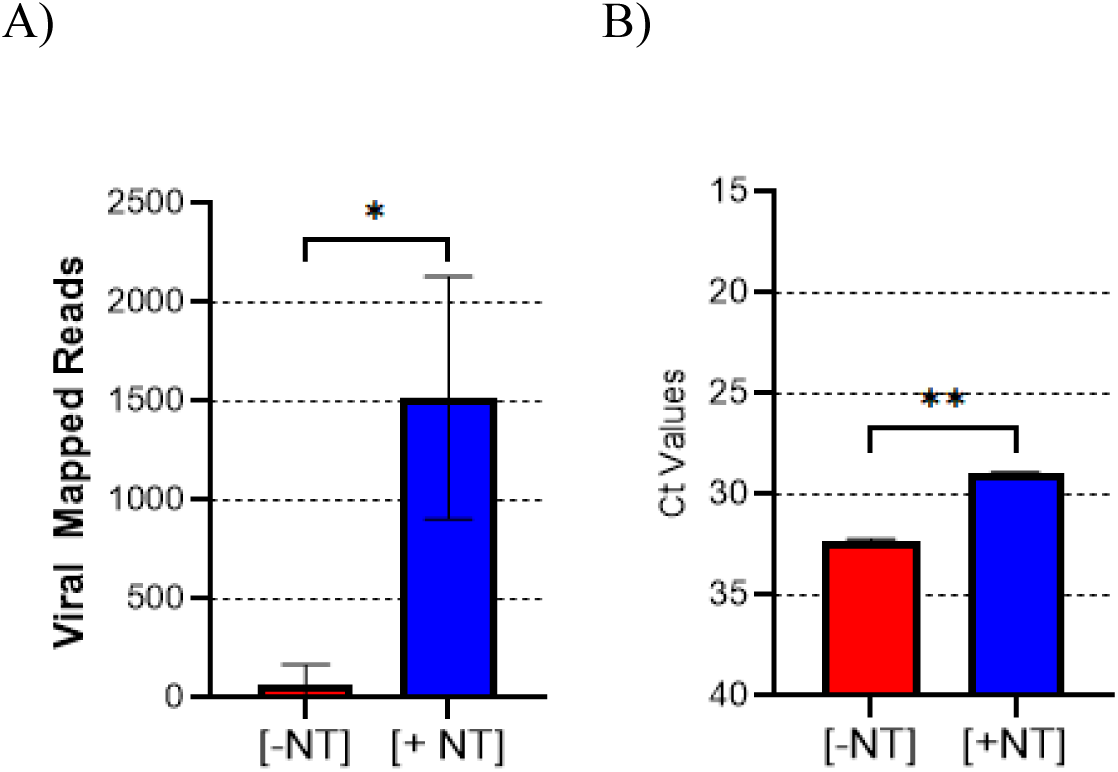
Nanotrap Particle Workflow 1 Improves Sequencing of Contrived Influenza A Samples. Heat-inactivated H1N1 was spiked into VTM at 10^6^ TCID_50_/mL and samples were processed using Nanotrap Particle Workflow 1 [+NT] or the RNEasy Kit alone [−NT]; n=3 for both processes. Samples then underwent sequencing on a ONT MinION R.9 flow cell (**A**) or RT-PCR (**B**). [+NT] were compared to [−NT] by paired *t*-test in order to assess significance of increased viral detection. * p<0.05, ** p<0.01.

**Fig. 6:**
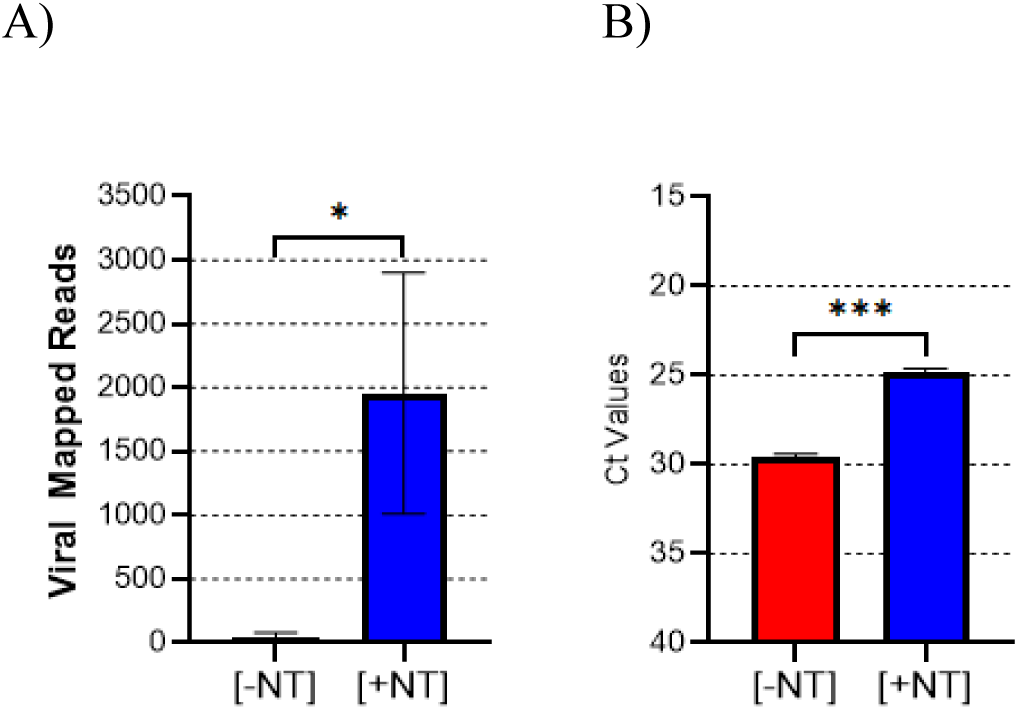
Nanotrap Particle Workflow 1 Improves Sequencing of Contrived RSV Samples. Heat-inactivated RSV was spiked into VTM at 10^6^ TCID_50_/mL and samples were processed using Nanotrap Particle Workflow 1 [+NT] or the RNEasy Kit alone [−NT]; n=3 for both processes. Samples then underwent sequencing on a ONT MinION R.9 flow cell (**A**) or RT-PCR (**B**). [+NT] were compared to [−NT] by paired *t*-test in order to assess significance of increased viral detection. * p<0.05, *** p<0.001.

## Discussion

Recent advancements in next-generation sequencing and post processing amplification techniques have decreased the viral titers required for successful sequencing runs and accurate mutation detection ^26,27^. While these advancements have generally improved the sensitivity of sequencing applications, for certain sequencing platforms there are still a significant portion of clinically relevant viral samples that cannot be sequenced due to low viral titers, samples in which there are insufficient nucleic acid molecules for bioinformatics tools to cover the entire reference genome while confidently distinguishing biological variation from error. More specifically, sequencing platforms “basecall”, or create readouts of the nucleotide fragments from the raw signals generated by the sequencer from the processed RNA samples. These basecalled fragments are compared and mapped against a database to identify the genomic taxonomy of the fragment. During this mapping process, differences will propagate between the analyzed sample and reference genome. These differences in readouts can be due either to real genetic mutations or to erroneous basecalling, the latter of which could be caused by either the sequencer itself or by insufficient nucleic acid material. Increasing the number of basecalled fragments clarifies this problematic overlap with greater read depths, increasing the confidence that detected variations are biological and not an artifact ^11–15^.

The Oxford Nanopore MinION sequencing platform is a compact, low complexity third-generation sequencing technology that has significantly reduced the upfront cost typically associated with sequencing. Although this technology has many appealing advantages, the usable clinical sample pool is restricted to higher titer samples due to sensitivity and accuracy limitations ^28–30^. We saw this limitation as an opportunity to examine potential enrichment strategies to enhance the amount of nucleic acid material being sequenced, increasing the available pool of clinical samples. To that end, we applied the Nanotrap particle front-end virus capture and concentration method to both contrived VTM and diagnostic remnant samples.

Nanotrap Particles significantly improved sequencing results by capturing and concentrating SARS-CoV-2 from contrived samples, improving the output of two standard RNA extraction methods. Furthermore, we identified a general working concentration range of SARS-CoV-2 in which Nanotrap particles were shown to significantly increase the viral mapped reads of the ONT Mk1C sequencing platform. Sequencing and RT-PCR improvements were seen for both Nanotrap Particle Workflows 1 and 2. Relative to the results delivered by the RNA extraction kits without Nanotrap particle pre-processing, both workflows significantly improved total viral mapped reads of SARS-Cov-2 at multiple concentrations. Of the two workflows, greater improvements were seen with the column-based Nanotrap Particle Workflow 1 over its comparator. This workflow employed a larger sample volume, allowing for a more significant amount of enrichment relative to the comparator.

For the magnetic bead-based RNA extraction kit approach, viral mapped reads were generally higher across all concentrations for samples processed with and without Nanotrap particles, relative to the column-based RNA extractions. This suggests greater RNA extraction efficiency of the magnetic bead extraction kit, binding and eluting a higher percentage of RNA. RT-PCR results supported this theory; we observed that the magnetic bead approach allowed detection of SARS-CoV-2 at a 10-fold lower concentration than the column-based approach. While viral mapped reads were higher for Nanotrap Particle Workflow 2 vis-a-vis Nanotrap Particle Workflow 1, the concentration range for which Nanotrap particles significantly improved the number of reads was similar for both workflows, improving sequencing results for the higher concentration samples. RT-PCR results showed Nanotrap particle enrichment was efficacious for lower concentration samples for both workflows, potentially suggesting that on an alternative sequencing platform with greater overall sensitivity, Nanotrap particles could also improve sequencing of these lower titer virus samples.

Additionally, results indicate that Nanotrap particle workflows improve sequencing and RT-PCR results of clinically relevant diagnostic remnant samples as compared to the workflows without Nanotrap particles. It appears that Nanotrap particles enhance remnant diagnostic sample sequencing results more significantly than in contrived VTM samples. We postulate that VTM collected from humans typically contains greater biological debris, and as a result, the workflows without Nanotrap particles are more likely to be impacted by inhibition while the Nanotrap particle pre-processing reduces this detrimental material through additional sample clean-up. It is possible that the Nanotrap particle architecture enables the capture of the viral material of interest while reducing host cell debris and other contaminating material. The sequencing library preparation workflow assessed here relied on a polymerase based amplification step which could be negatively impacted by human cellular material, cleaning up background material while capturing and concentrating viral material would allow the Nanotrap workflow to improve this amplification step even further generating greater total viral mapped reads for diagnostic remnant VTM samples.

Nanotrap particles significantly improved viral mapped reads of a majority of the diagnostic remnant samples for both workflows. As a result, viral genome coverage also increased for a majority of diagnostic remnant samples, increasing by 80% in certain diagnostic remnant samples. If these diagnostic remnant samples represented a larger pool of clinical samples, this data suggests Nanotrap particles would have significantly increased the fraction of samples that could be used for sequencing. In samples where there was not a statistically significant improvement, the samples processed without Nanotrap particles were already generating relatively complete genome coverage. Data generated using the diagnostic remnant samples also appeared to suggest that greater mapped reads are required to obtain complete coverage of the SARS-CoV-2 genome when using a magnetic-bead-based workflow as opposed to the column workflow. Although this did not seem to reduce the utility of the magnetic-bead based workflow as we observed that samples with the same viral titer typically generated more total viral reads when using the magnetic-bead-based extraction kit. These differences between the extraction kits could be due to the column having a higher affinity for larger viral RNA fragments while having a lower overall extraction efficiency, allowing for smaller amounts of RNA to more completely cover the SARS-Cov-2 genome, while the magnetic-bead-based RNA extraction inputs more but generally smaller RNA fragments.

In order for sequencing to become more useful in public health settings, sample throughput and workflow considerations must be addressed. Automated systems, such as the KingFisher Apex System, are readily scalable and already used as a processing tool in the clinical laboratories ^31^. Nanotrap particles improved the sequencing results of ten positive diagnostic remnant samples when processed using Workflow 3, demonstrating utility in a high throughput automated system. It is worth noting that this automated method would enable the processing of 96 samples in 1 hour, which is significantly faster and far more user friendly than the manual-column extraction method, making this an attractive proposition for medium-to high-throughput laboratories.

In addition to SARS-CoV-2, Nanotrap particles have demonstrated use in capturing a broad range of viruses, including respiratory pathogens ^17–21^. However, to date, no viral sequencing data has ever been published when using a Nanotrap particle workflow. Here, we confirmed previous reports showing Nanotrap particles can also capture and enrich both Influenza A and RSV, two common respiratory viruses. Demonstrating that our Nanotrap workflow is compatible for sequencing of multiple respiratory pathogens, increasing viral mapped reads of both viruses. This suggests that Nanotrap particle workflows can be used for the improvement of broad-scale viral detection by sequencing.

This study contains certain limitations, beginning with the number of samples assessed. Sufficient numbers of replicates were tested to determine a positive improvement provided by the Nanotrap particle process when sequencing VTM samples, but more samples should be run to better and more accurately quantify the fold-enrichment the workflow can provide. We also did not directly assess the Nanotrap particles’ ability to improve sequencing of different SARS-CoV-2 variants. Given the general viral capture nature of the Nanotrap particles, we expect that sequencing improvements seen with the wild-type SARS-CoV-2 would correspond to improvements across most other SARS-CoV-2 variants. Future experiments could test this hypothesis using specific known variant samples, assessing Nanotrap particle enrichment on the basis of increased variant detection. We could create an artificial testing pool of diagnostic remnant samples with known variants at lower titres to examine if the Nanotrap particle workflow can improve the number of available clinical samples for sequencing. We also only examined contrived influenza A and RSV samples, so we cannot definitively say at this time that the current Nanotrap particle process is capable of sequencing these respiratory pathogens in more biologically complex media. We also did not examine what occurs in samples that are co-infected with multiple viruses. This could potentially bias the Nanotrap particles towards a specific virus should that virus have higher affinity to the Nanotrap particles than others. Future experiments would examine how Nanotrap particles behave in a co-infected sample, along with using more clinically relevant diagnostic remnant samples containing influenza A or RSV.

Overall, this study indicates that Nanotrap particle enrichment allows for sequencing of lower titer clinical samples in VTM using the ONT MinION Sequencer, which otherwise may not have been suitable for sequencing. Because our method requires no filtration or centrifugation steps, this approach is compatible with medium- and high-throughput environments, including the KingFisher Apex platform. Additionally a Nanotrap particle concentrating method paired with a ONT sequencing platform allows for a powerful sequencing tool that could potentially be deployed in an area with lack of access to more traditional sequencing or sample processing equipment.

There is room to explore additional applications of this approach to alternative sample types, including oral fluid (which could be used for less invasive viral respiratory testing) and wastewater (which could be used to conduct surveillance of viral respiratory pathogens in communities). Going forward, we plan to address each of these areas so that we can continue examining viral surveillance applications with this versatile enrichment technology.

## Acknowledgements

This publication was supported by the SBIR Grant 1 R43IP001142-01 from the Centers for Disease Control and Prevention and by the NIH Rapid Acceleration of Diagnostics (RADxSM) initiative funded in whole or in part with Federal funds from the National Institute of Biomedical Imaging and Bioengineering, National Institutes of Health, Department of Health and Human Services, under Contract No. 75N92020C00021. Its contents are solely the responsibility of the authors and do not necessarily represent the official views of the Centers for Disease Control and Prevention or those of the National Institutes of Health.

## Author Contributions

P.A, N.S., D. G., J. F.,and K.B. carried out the experiments, contributing to viral capture, viral extraction, and viral detection. A.P., D.M., S.M., S.S.,R.K.,C.R.,S.M,D.W.,J.S.,F.N., O.S., A.M.,K.K., and K.H. contributed to the production and quality control of the Nanotrap® particles. P.A.,R.B., T.J., B.L.,S.B.,R.D., and R.A.B. contributed to data presentation and formatting. P.A., S.B., R.A.B., and B.L. analyzed, interpreted the data, and were involved in the experimental design. P.A.,R.B., T.J., B.L.,S.B.,R.D., and R.A.B. were involved in the writing and editing of the manuscript while also providing the overall direction and coordination of this study. All authors approved the manuscript

